# An Allosteric Model for Electromechanical Coupling in Cardiac CNBD Channels

**DOI:** 10.1101/2025.07.28.666606

**Authors:** Gucan Dai

## Abstract

Ion channels in the cyclic nucleotide-binding domain (CNBD) family, including hyperpolarization-activated cyclic nucleotide-gated (HCN) channels and human ether-à-go-go-related gene (hERG) channels, play pivotal roles in regulating cardiac action potentials. HCN channels are uniquely activated by hyperpolarization, rather than depolarization, a critical mechanism for controlling the involuntary pacemaker activity of the heart. In contrast, hERG channels are depolarization-activated and mediate K^+^ currents essential for action potential repolarization. Notably, certain hERG mutations, including those associated with long-QT syndrome, can induce biphasic activation by both hyperpolarization and depolarization. Despite the diverse voltage-dependent gating behaviors observed in CNBD channels, a unified mechanistic framework remains lacking. Here, we propose an allosteric model for their electromechanical coupling, featuring a single voltage-sensor transition coupled to two distinct conformational coupling modes between voltage-sensing and pore domains. With only three or four free parameters, this model recapitulates the biphasic U-shaped and bell-shaped conductance-voltage relationships commonly seen in CNBD channels. Fluorescence anisotropy-based homo-FRET experiments employing site-specifically incorporated noncanonical amino acids provide further support for the hypothesis, suggesting that the S5 helix movement plays a key role in hyperpolarization-dependent activation, while S4-S6 helix interactions are required for depolarization-dependent gating.

## Introduction

Hyperpolarization-activated cyclic nucleotide-gated (HCN) ion channels and human ether-à-go-go (EAG) channels, also known as KCNH voltage-gated potassium channels (K_v_ 10-12), belong to the cyclic nucleotide-binding domain (CNBD) family within the superfamily of voltage-gated ion channels (VGICs) (1, 2). They are pivotal in determining the rhythm of cardiac action potentials. HCN channels become active in response to membrane hyperpolarization, a critical process for regulating the pacemaker activity of sinoatrial-node cells in the heart. This mechanism is not only critical for normal cardiac pacemaking but also holds relevance for understanding cardiac arrhythmias such as bradycardia and atrial fibrillation (1, 3-6). This type of hyperpolarization-dependent channel opening is contrasted with most other VGICs, including KCNH channels, which are activated by membrane depolarization (1, 7). For HCN channels, second messenger cyclic nucleotides bind to the CNBD directly and potentiate HCN channel activity (8, 9). This modulation of the main cardiac isoform HCN4 by cAMP forms the basis for sympathetic control over heart rate (3, 10). Furthermore, the human ether-à-go-go-related gene (hERG, K_v_11) channel, member of the KCNH subfamily underlies the rapid delayed rectifier potassium current (I_kr_), which is important for the repolarization phase of the cardiac action potential (11-13).

Despite their functional diversity, CNBD family channels share a conserved architecture composed of four similar or identical subunits arranged around a central pore. Each subunit comprises essential components, including a voltage-sensing domain (VSD) consisting of four transmembrane helices (S1 to S4), a pore domain (PD) formed by S5-S6 transmembrane helices that constitute the channel gate, a selectivity filter, a C-terminal C-linker domain, and a cyclic nucleotide-binding (or homology) domain (CNBD) (2, 5, 14) (Fig. 1). The S4 helix contains positively charged arginines that directly sense transmembrane voltage. The voltage sensor S4 helix moves in the same direction for both HCN and hERG channels, suggesting a reversed coupling between the VSD and the PD that underlies the opposite gating polarities (15-19). Like HCN channels, hERG channels contain a VSD and a PD. The intracellular parts of each subunit include an N-terminal eag domain and a C-terminal cyclic nucleotide-binding homology domain (CNBHD) (13, 20-27). The CNBHD in hERG is insensitive to cyclic nucleotides like cAMP and cGMP (21, 28, 29). Notably, specific mutations or cross-linking strategies could reverse the gating polarity of HCN channels, causing them to activate upon depolarization (18, 30). Additionally, engineered or mutated CNBD channels may exhibit biphasic gating behavior, characterized by U-shaped or bell-shaped conductance-voltage (G-V) relationships (31-34). Despite their structural similarities, it remains unclear how these channels give rise to such diverse voltage-dependent behaviors.

**Fig. 1.**
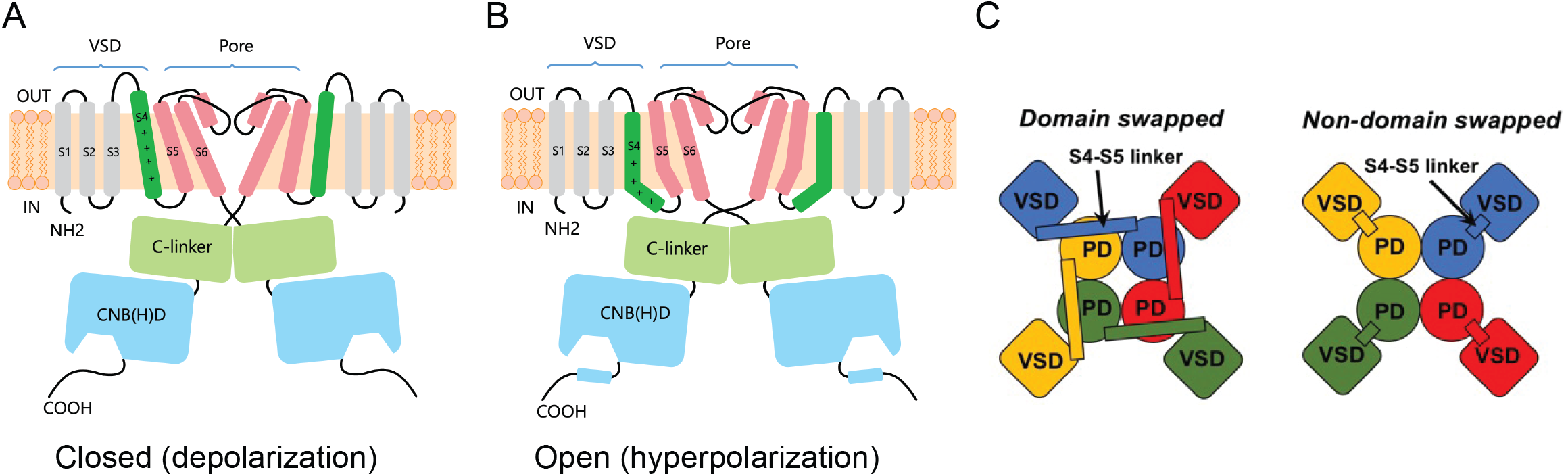
Key Structural Features of CNBD family channels. (**A-B**) Basic shared architecture of CNBD channels showing transmembrane domains and C-terminal C-linker and CNB(H)D domains. The divergent N-terminal regions are not shown. Particularly, for the unique hyperpolarization-dependent activation in HCN channels, the downward movement of the S4 likely tilts the S5 helix, which rearranges the S6 pore helix, leading to channel opening (B). (**C**) The comparison of non-domain-swapped VSD-PD configuration in CNBD channels compared to the domain-swapped configuration seen in classic VGICs.

CNBD family channels are macromolecules with multiple functional modules or domains. An unexpected finding was that all CNBD channel structures showed a remarkably different tetrameric assembly called non-domain-swapped VSD-PD configuration compared to the domain-swapped configuration seen in classic VGICs (Fig. 1C) (24, 35-39). In contrast to the domain-swapped VSD/PD assembly such as for voltage-gated Na^+^ and Ca^2+^ channels, the VSD and PD of the same subunit are close to each other in CNBD channels (35, 39, 40). An intriguing and perhaps unique aspect of CNBD channel assembly is their ability to form functional channels from split constructs, in which the VSD and PD are expressed as separate, covalently disconnected polypeptides (31, 41). The VSD-PD coupling of non-domain-swapped CNBD channels differs from the canonical electromechanical coupling in domain-swapped channels (31, 41, 42), and likely entails more than one coupling mode (17, 43).

In this study, we present an allosteric model for CNBD-family channels in which gating is governed by one or two electromechanical coupling pathways. Guided by computational modeling and fluorescence anisotropy, we propose that voltage-dependent interactions among S4, S5, and S6 helices cooperatively transduce voltage sensing into pore opening. Our model demonstrates how dual voltage-dependent coupling factors enable functional diversity across CNBD channels. Fluorescence anisotropy results further support the presence of distinct helix rearrangements underlying channel activation in HCN and hERG channels, mediated by S4-S5 or S4-S6 interactions, offering a mechanistic framework that reconciles previous structural and functional observations. These findings redefine the paradigm of voltage-dependent gating in CNBD channels, highlighting voltage-dependent coupling as a key determinant of their allosteric activation.

## Results

### A modular allosteric model for CNBD channel gating

This hypothesis-driven work proposes a new gating model to explain the diverse voltage-dependent gating polarities observed in the CNBD channel family. Here we present a modular allosteric framework to describe a voltage-gated ion channel with VSD modules, each capable of bidirectional coupling to the pore module via distinct modes: facilitation or positive coupling factor D, and inhibition or negative coupling factor H (Fig. 2A). In other words, D increases with depolarization whereas H increases with hyperpolarization. To simplify the model, we assume that four identical VSDs act concertedly and must be simultaneously activated to gate the pore. Thus, the coupling factors are raised to the 4^th^ power to represent maximal cooperativity. The modular gating scheme can be computed for both positive and negative voltage-dependent regulation using a cubic (8-state) model (Fig. 2B). The open probability reflects three key contributions: baseline spontaneous pore opening, opening by single coupling mode between VSD and PD, and opening by dual coupling modes between VSD and PD. This formulation captures mixed cooperativity, where H and D compete to modulate the pore, enabling fits to various gating behaviors effectively. Unlike traditional modular gating models that assume voltage-independent coupling, this approach uniquely accommodates a diversity of gating polarities.

**Fig. 2.**
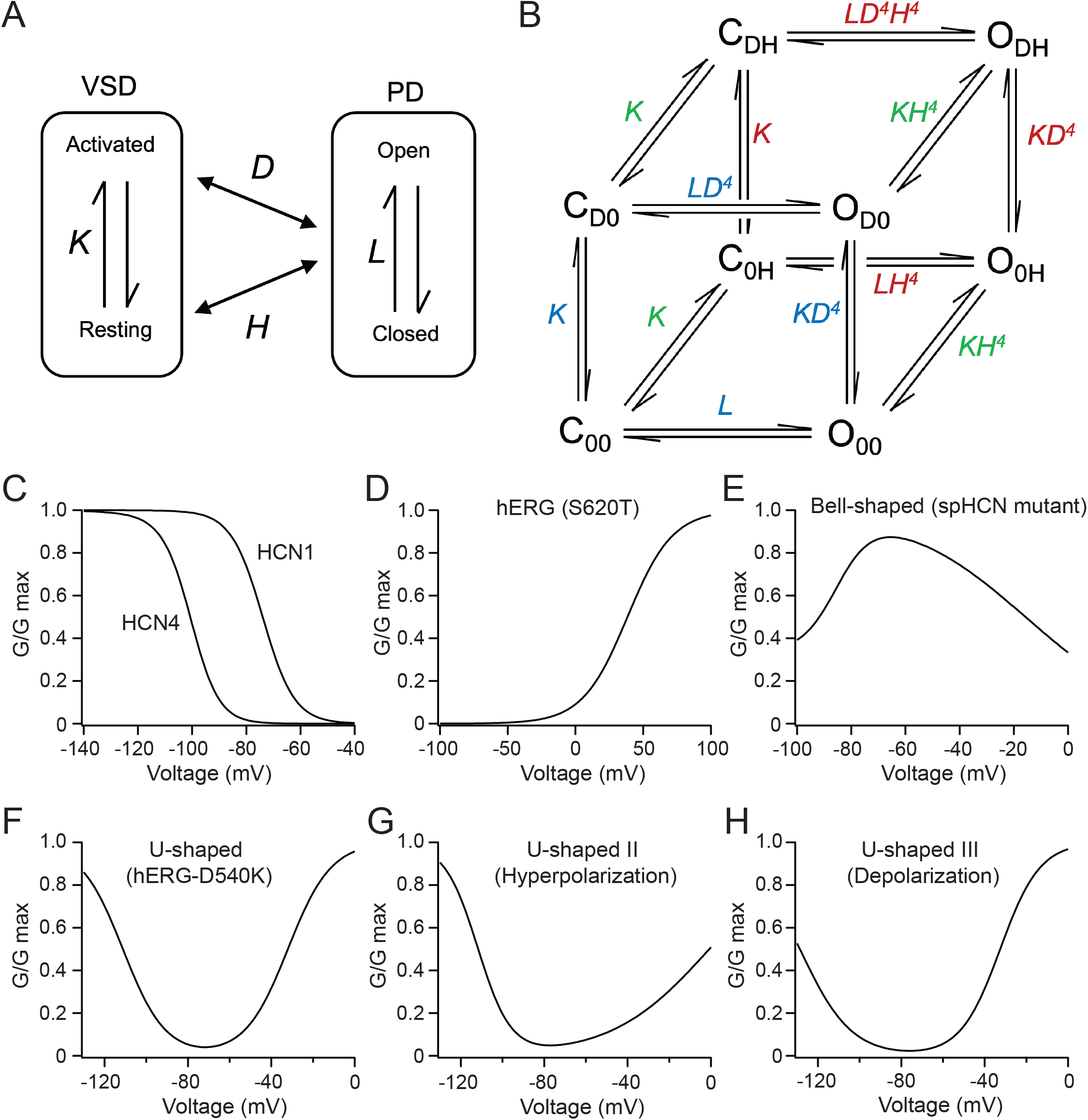
An allosteric model for bidirectional voltage-dependent gating in CNBD channels. (**A**) Schematic of the dual-coupling modular framework, where the voltage-sensing domain (VSD) bidirectionally regulates the pore domain (PD) via facilitation (D, positive coupling) or inhibition (H, negative coupling). (**B**) Cubic 8-state gating scheme integrating baseline pore opening, single-coupling (D or H), and dual-coupling (D and H) contributions. The model includes four closed states (C_00_, C_D0_, C_DH_, C_0H_), representing the channel’s closed, pre-open conformations under different coupling conditions: no coupling C_00_, D-coupling alone (C_D0_), both D- and H-coupling (C_DH_), or H-coupling alone (C_0H_). The corresponding open states (O_00_, O_D0_, O_DH_, O_0H_) mirror these configurations, reflecting pore opening under equivalent coupling conditions. The model assumes four identical VSDs acting cooperatively. (**C**-**D**) Simulated conductance-voltage (G-V) relationships for HCN1, HCN4, and hERG channels, fitted using the dual-coupling model (parameters in Table S1). Data points were sampled every 1 mV. HCN1 and HCN4 exhibit hyperpolarization-driven opening, while hERG shows hyperpolarization-driven closure. (**E**- **H**) Simulated biphasic G-V curves: bell-shaped (E, e.g., spHCN mutants) and U-shaped (F, e.g., hERG- D540K), using the same allosteric model. The model captures diverse gating polarities, including hyperpolarization-shifted (U-shape II) in G, and depolarization-shifted (U-shape III) profiles in H.

To validate the effectiveness of this model, we successfully simulated the gating profiles of HCN1, HCN4, and hERG channels (Fig. 2 C-H, and Table 1). Regarding the two coupling factors, only H is active in HCN channel opening, whereas only D is active in hERG channel opening. Key differences in HCN and hERG gating parameters emerge in two aspects: first, HCN channels require minimal spontaneous opening (small L values), while hERG channels exhibit significantly larger L, reflecting their greater intrinsic open probability (24). This indicates that the VSD presents a strong inhibitory effect to the pore in HCN channels, ensuring channel closure at depolarized voltages (31). Second, HCN1 and HCN4 (z = 3) require larger gating charges than hERG (z = 2), consistent with hERG’s shallower activation voltage dependence (13, 44) (Table 1). The greater voltage-dependent movement of the S4 helix in HCN channels, compared to hERG, reflects the involvement of more voltage-sensing arginines traversing the membrane electric field during activation (15, 16, 44-46).

**Table 1.**
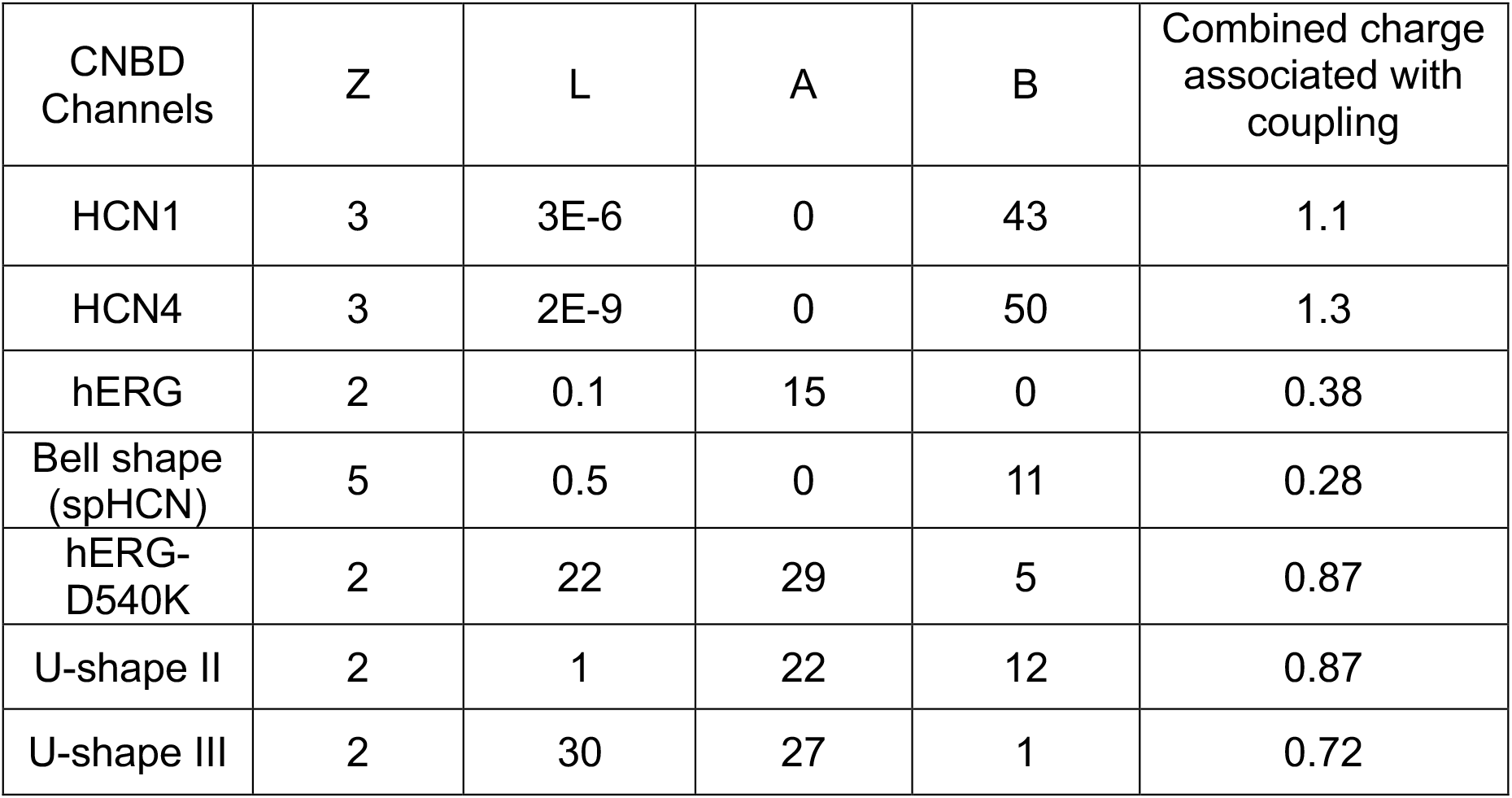
List of the four free parameters and the combined charge associated with coupling factors for the 8-state modular gating scheme simulating various CNBD-channel gating features.

| CNBD Channels | Z | L | A | B | Combined charge associated with coupling |
| --- | --- | --- | --- | --- | --- |
| HCN1 | 3 | 3E-6 | 0 | 43 | 1.1 |
| HCN4 | 3 | 2E-9 | 0 | 50 | 1.3 |
| hERG | 2 | 0.1 | 15 | 0 | 0.38 |
| Bell shape (spHCN) | 5 | 0.5 | 0 | 11 | 0.28 |
| hERG-D540K | 2 | 22 | 29 | 5 | 0.87 |
| U-shape II | 2 | 1 | 22 | 12 | 0.87 |
| U-shape III | 2 | 30 | 27 | 1 | 0.72 |

### Simulating bipolar voltage-dependent gating of CNBD channels

We also simulated the biphasic gating, mirroring features observed in several mutant or engineered CNBD channels that exhibit either bell-shaped or U-shaped G-V relationships. The bell-shaped G-V relationship was observed in sea urchin HCN channels (spHCN) following alanine substitutions at the C-terminal end of the S4 helix or a single R530A mutation within the S4 helix, all in the absence of cyclic nucleotides (31). We found that, in comparison to wild-type HCN channels, a positive coupling factor D was needed to simulate the bell-shaped G-V (Table 1). In this case, the D acts like an inactivation factor, and a similar strategy can be used to simulate the bell-shaped G-V in N-terminal PAS cap-truncated Eag channels with just one coupling factor (34). Additionally, an increased gating charge (z = 5, compared to z = 3 for wild-type hHCN1 and hHCN4) for the mutant spHCN-A6 channel was accompanied with the bell-shaped G-V, suggesting the scale of the voltage sensor S4 helix movement may be greater (Table 1) (31). Moreover, a symmetrical U-shaped G-V was seen in the case of hERG mutant D540K, and various additional mutations could produce U-shape II or III gating behaviors in hERG-D540K channels (32). The U-shaped G-V relationship was also observed in chimeric channels containing structural elements from HCN1 and Eag channels as well as in truncated Eag channels (33, 34). We found that reproducing U-shaped G-V relationships—whether symmetrical, hyperpolarization-dominant (U-shape II), or depolarization-dominant (U-shape III)—often requires both coupling factors D and H. However, the characteristic U shape can still emerge with coupling factor D alone. Incorporating both D and H increases the model’s flexibility, enabling it to capture a broader range of bell-shaped or U-shaped gating behaviors. Moreover, one common feature for the bipolar G-V exhibiting channels is that a larger L is needed, indicating a greater intrinsic open probability.

As an unconventional approach to model electromechanical coupling—where coupling factors are allowed to vary with voltage—we calculated the combined charge associated with the H and D coupling factors for each simulated gating behavior of CNBD channels (Table 1). These values are small (< 1) compared to the gating charge contributed by the primary S4 movement in hERG and in all biphasic gating behaviors. The coupling-associated charges are 1.1 for HCN1 and 1.3 for HCN4 (Table 1), which may reflect the motion of arginine residues in the extracellular portion of the extra-long S4 helix during the coupling process (47, 48). These estimates suggest that, during this noncanonical coupling between the voltage sensor and the pore domain, subtle rearrangements of the S4 helix—or changes in its interactions—could generate small but noticeable charge movements within the membrane electrical field.

### Alternative allosteric models using a similar modular scheme

We also evaluated simpler allosteric schemes with four or six states (Fig. 3). The four-state model assumes that all subunits are equivalent and each couples to the pore through both D and H. In contrast, the six-state model assumes that one subunit couples to the pore through either D or H, but not both. Although both models reproduced the wild-type features of HCN and hERG channels and several features of biphasic G-V relationships, the four-state model failed to capture the characteristic U-shape II behavior, which has higher open probability at hyperpolarized voltages than at depolarized voltages. Nonetheless, this much simpler model can largely reproduce the wild-type and biphasic gating behaviors of CNBD channels. In contrast, the six-state model fails to recapitulate any U-shaped gating features (Fig. 3). These limitations indicate that the more complex (eight-state) cubic model may be necessary to account for the full diversity of CNBD-channel gating. Our results further suggest that the four CNBD channel subunits can couple to the pore in subtly distinct ways within the same channel, likely shaped by local lipid environments and modulatory intracellular domains (2, 19, 49).

**Fig. 3.**
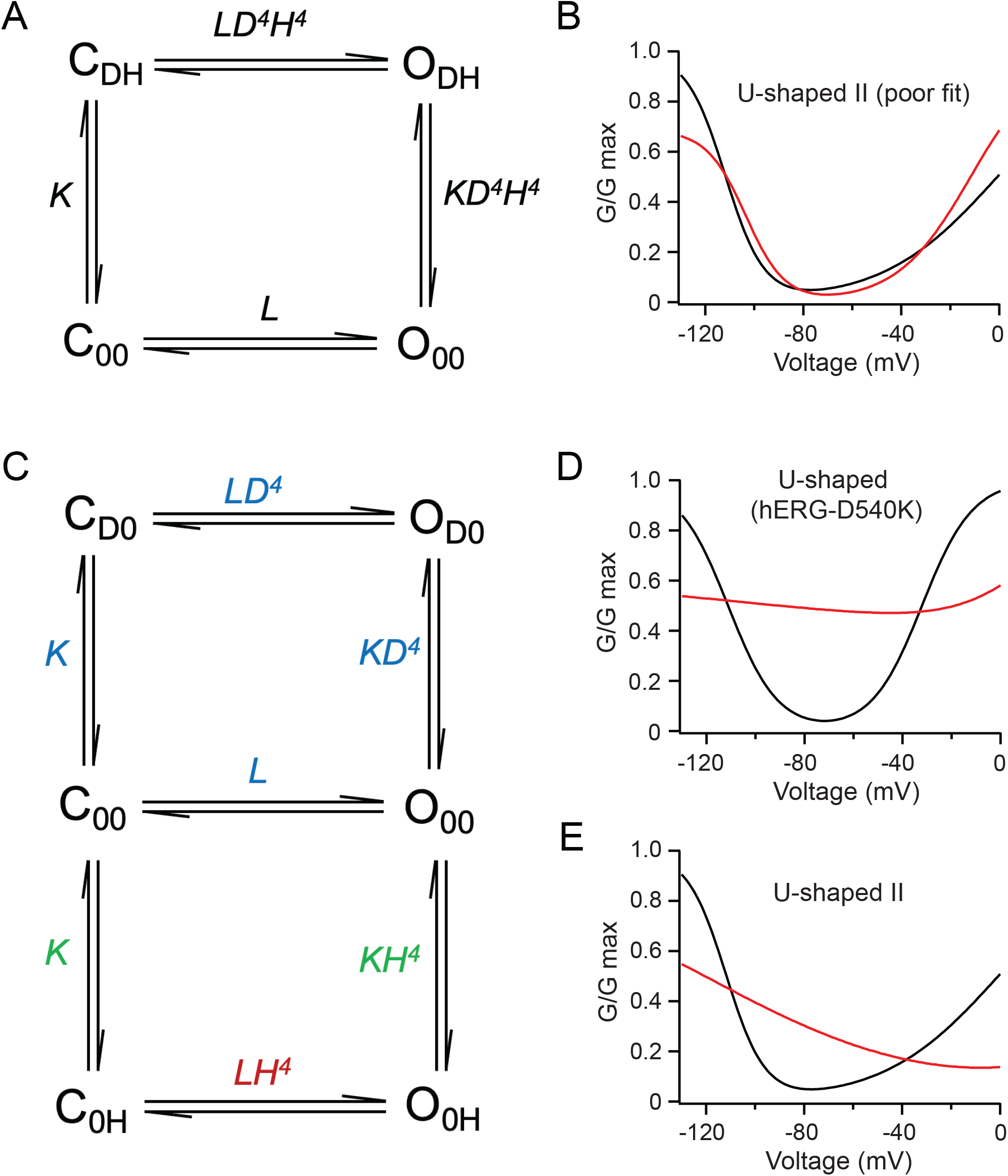
Alternative 4-state and 6-state models were unable to fully recapitulate the observed CNBD gating behaviors. (**A**) Scheme of the four-state model where all subunits are equivalent and each couples to the pore through both the D and H couplings. (**B**) The 4-state model can fit most of the gating behaviors of CNBD channels well except for the hyperpolarization-favored U-shaped gating (type II). In this case, the parameters of the fit (red curve) are z = 3.17, L = 2.18, A = 27, B = 10.4. (**C**) The 6-state model that assumes that the S4 can only be coupled to the pore through either the D or the H coupling. (**D-E**) the 6-state model failed to fit all the U-shaped gating behaviors. Two examples are shown for this model. The free parameters of the red curve in D (U-shaped hERG-D540K) are z = 2, L = 1.4, A = 13.5, B = 1. The free parameters of the red curve in E (U-shaped type II) are z = 2, L = 0.16, A = 8.1, B = 5.3. The K_0_ = 1.26E+6 for all models in this figure.

### Homo-FRET experiments support the key role of S5 helix in electromechanical coupling for HCN

To investigate structural mechanisms underlying the dual coupling modes suggested by our allosteric model, we focused on the S5 helix—a known gating element that relays the S4 movement to open the pore in HCN channels (17, 47, 50). Using stop-codon suppression to incorporate fluorescent noncanonical amino acids, we employed fluorescence anisotropy-based homo-FRET to gain sights into the lateral movements of the S5 helix in hHCN4 and hERG channels. To site-specifically label the channels, we used the established amber stop-codon suppression strategy to incorporate the noncanonical amino acid L-Anap to the V419 site within the S5 helix of hHCN4 and the equivalent F551 site of hERG (Fig. 4 A-C) (51-53). We also engineered channel constructs for a comparison of the C-linker movement for hHCN4 and hERG channels by incorporating L-Anap to the Q536 and L678 sites, within the A’ helix of the hHCN4 and hERG channels, respectively (Fig. 4 A-C).

**Fig. 4.**
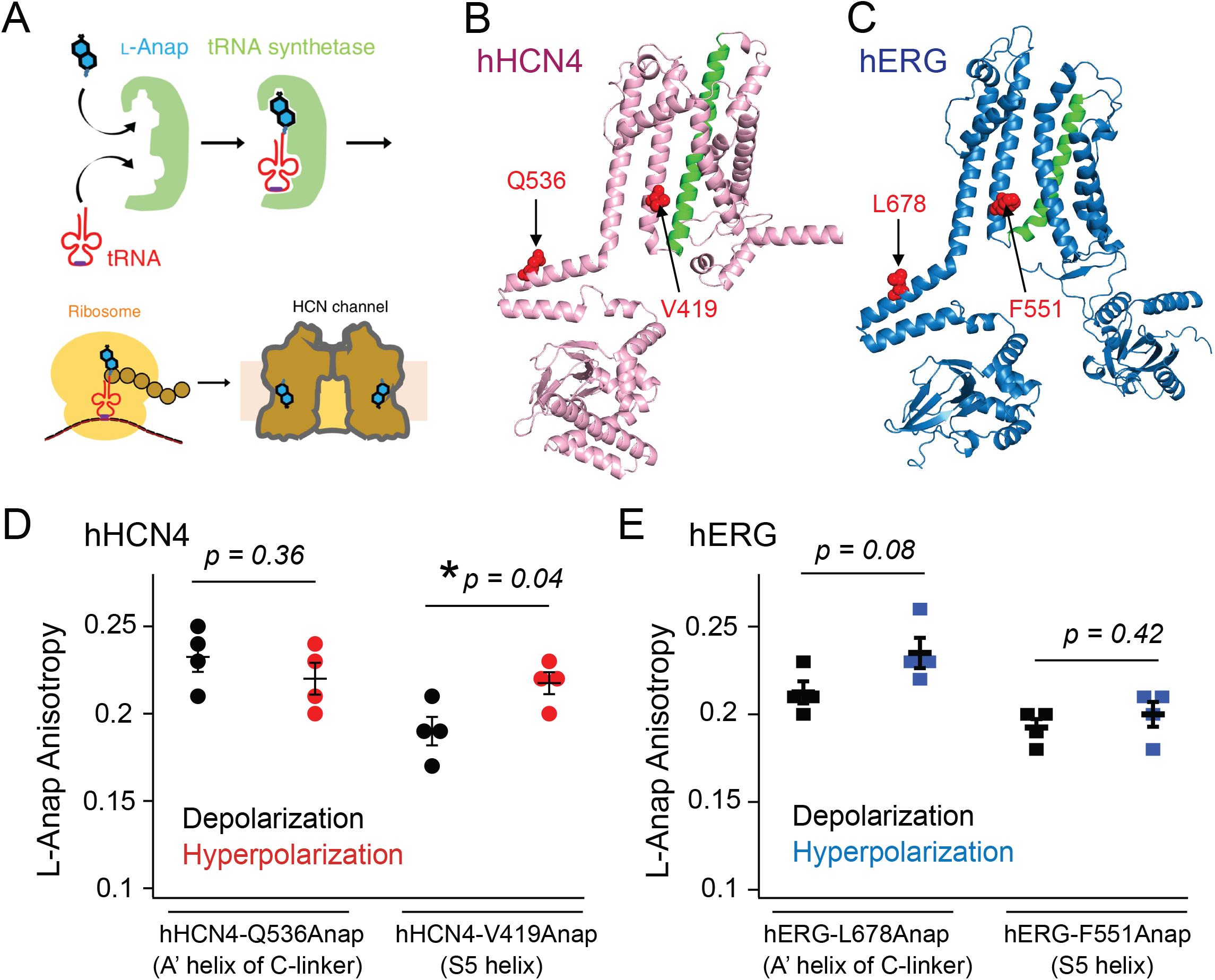
Homo-FRET suggests S5 helix movement for channel opening, unique in HCN channels. (**A**) The amber stop-codon suppression strategy for incorporating L-Anap. (**B**-**C**) Schematic of hHCN4 and hERG channels showing L-Anap labeling sites in S5 (V419/F551) and C-linker (Q536/L678). The structures shown are single subunits from either AlphaFold generated models of hHCN4 or hERG. (**D**-**E**) Fluorescence anisotropy traces during hyperpolarization (NMDG) and depolarization (KCl). In D, hHCN4-V419Anap (S5) shows increased anisotropy (reduced homo-FRET) during hyperpolarization, indicating S5 movement, while Q536Anap (C-linker) remains unchanged. In E, hERG-F551Anap (S5) exhibits no anisotropy shift, whereas L678Anap (C-linker) shows modest changes. Data suggest S5 movement drives HCN activation but is dispensable for hERG gating, supporting distinct coupling mechanisms. Error bars represent SEM, n = 4 for all conditions, two-sided t-test was used.

Upon excitation with polarized light, fluorescence anisotropy measures the polarization of emission from L-Anap incorporated within hHCN4 and hERG channels, expressed in tsA-201 cells. This allows the quantification of homo-FRET by detecting depolarization resulting from energy transfer between nearby fluorophores. A decrease in anisotropy indicates increased homo-FRET efficiency, reflecting a closer proximity between L-Anap molecules (54). Homo-FRET-based anisotropy is particularly sensitive to the distance between fluorophores. One advantage is that it is less affected by factors such as fluorophore photobleaching, making it a robust tool for studying molecular movement over time (55). Although we cannot distinguish between homo-FRET occurring between adjacent or diagonal subunits, this design offers an efficient approach to assess the lateral movement of backbone helices within a tetrameric channel. Additionally, rotational depolarization arising from local side-chain motion is expected to be largely saturated for small amino acids such as L-Anap, whose local rotational correlation times (θ) are typically in the sub-nanosecond (picosecond) regime, substantially shorter than the fluorescence lifetime (τ) of L-Anap (on the order of a few nanoseconds). Under these conditions (θ ≪ τ), based on the Perrin equation, even substantial changes in side-chain mobility would be predicted to produce only modest changes in anisotropy, unlikely to fully explain any large effects we observed.

Distinct changes in Anap anisotropy were observed at specific sites within the hHCN4 and hERG channels under hyperpolarized and depolarized conditions, providing insights into the structural dynamics of these channels. Hyperpolarization was induced by an external solution containing 107 mM NMDG, while depolarization was achieved with 120 mM KCl. Under depolarization conditions, the measured anisotropy is generally higher for A′ helix sites compared to S5 helix sites, consistent with greater inter-subunit distances involving the A′ helix than those involving the S5 helix (Fig. 4 D and E). The results revealed no hyperpolarization-induced change in anisotropy for Q536Anap, suggesting that this site did not undergo significant conformational rearrangements during the voltage changes (Fig. 4D). However, a significant increase in anisotropy (a decrease in homo-FRET) for V419Anap during hyperpolarization indicated that the distance between fluorophores at this site within the hHCN4 S5 helix became more distant when at hyperpolarized voltages (Fig. 4D). Moderate changes for hERG-L678Anap and no change for hERG F551Anap imply that these regions remain relatively stable under the conditions tested (Fig. 4E). Overall, the most significant change in anisotropy was observed at V419 in the hHCN4 S5 helix, suggesting that this region may play a critical role in HCN channel activation in response to hyperpolarization. In contrast, the opening of KCNH channels is likely accompanied by the C-linker movement (37), consistent with the noticeable anisotropy change seen at the L678Anap site. Conversely, the lack of anisotropy change at the F551Anap site indicates that S5-helix movement may not be essential for hERG channel gating, aligning with the idea of distinct electromechanical coupling mechanisms between these channels.

### A hypothesis involving two coupling pathways for CNBD channel gating

This paper emphasizes the need for a new model for the EM coupling of CNBD channels, separate from the canonical model in domain-swapped VGICs. We propose that there are two coupling modes that control the voltage-dependent gating polarity for CNBD channels. Specifically, at hyperpolarizing voltages, the interaction between the S4 and the S5 helix promotes channel opening whereas the interaction between the S4 helix and the S6 helix prevents channel opening (Fig. 5). A mixture of these two modes of action leads to biphasic U-shaped G-V relationships, often observed in mutant CNBD channels (32-34, 56). Unlike conventional electromechanical coupling mechanisms in domain-swapped VGICs, we propose that CNBD-family channels rely on two distinct interactions: (i) H coupling, mediated by the S4-S5 helix interaction, and (ii) D coupling, involving direct interaction between the S4 and S6 helices. In HCN channels, the elongated S4 helix is positioned far from S6, whereas in hERG channels, the C-terminus of S4 interacts directly with S6, eliminating the need for S5 helix-mediated electromechanical coupling (Fig. 5). Although experimentally validating these dual coupling pathways remains technically challenging, our allosteric model supports the plausibility of this mechanism.

**Fig. 5.**
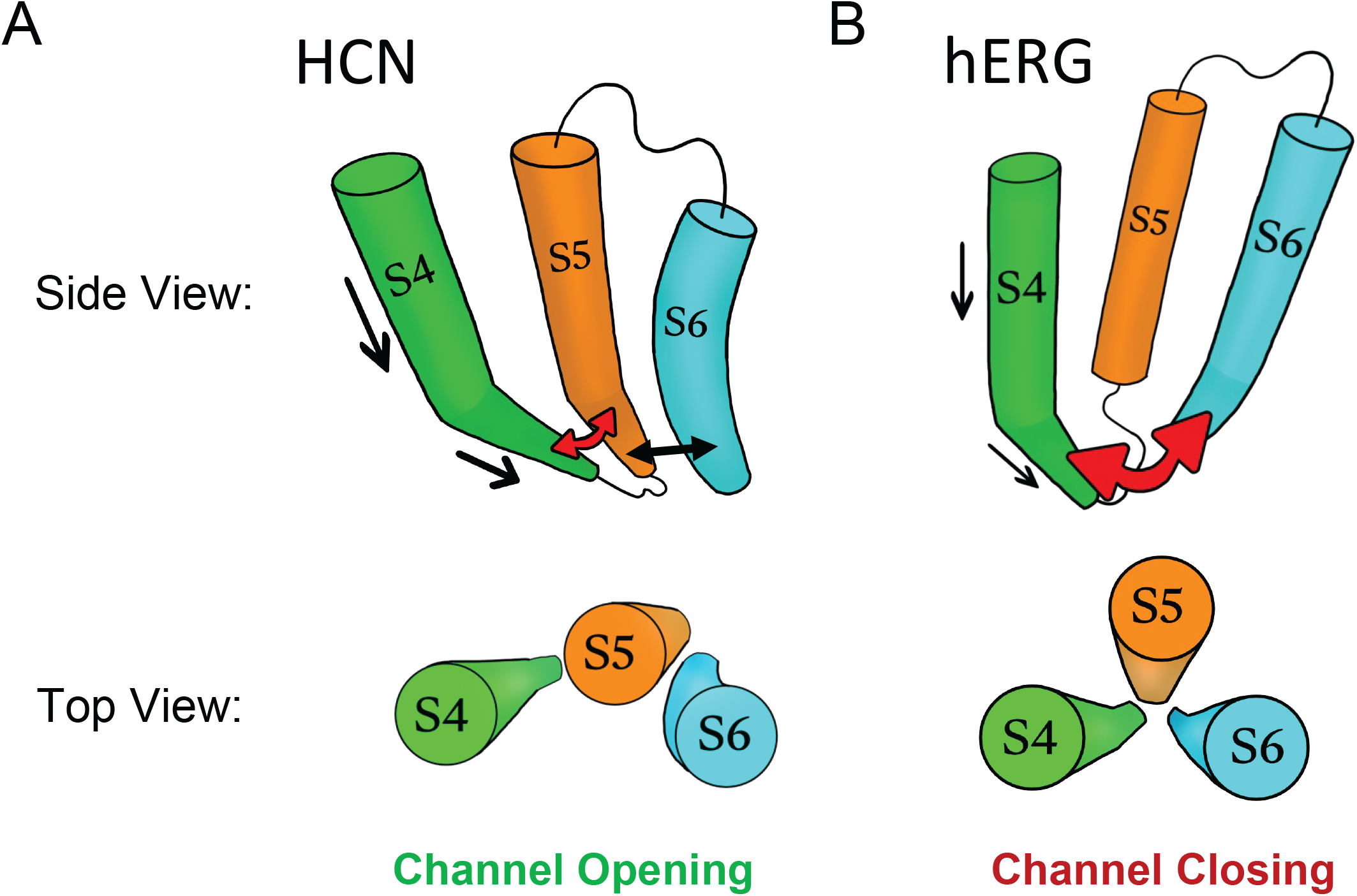
Cartoons highlighting the two electromechanical coupling modes for CNBD channels. (**A**) In HCN channels, downward movement and tilting of the S4 voltage sensor displaces the S5 helix, which then shifts the S6 helix to open the pore. S4 does not directly interact with S6. (**B**) In the S4-down conformation of hERG channels, S4 directly pushes against the S6 helix, forcing pore closure. S4 upward movement opens the pore. The cartoons were based on solved cryo-EM structures of HCN and hERG.

## Discussion

Our simple 8-state cubic gating model with four free parameters provides a possibility for describing the complex voltage-dependent gating in CNBD channels. Unlike classical sequential models, this framework accounts for multiple intermediate conformations, enabling it to capture bidirectional voltage sensitivity (both depolarization- and hyperpolarization-dependent gating) observed in different family members. It can explain how subtle differences in the VSD-PD coupling can yield divergent gating behaviors, optimizing channels for distinct roles in cardiac pacemaking, cAMP signaling, or neuronal excitability. Key innovations include the incorporation of dual voltage-dependent VSD-pore interactions and representation of intermediate states, offering testable predictions for future mutagenesis and electrophysiology studies. This approach not only unifies disparate gating phenotypes but also provides a mechanistic basis for disease-associated mutations that disrupt these transitions.

This allosteric model highlights several key features that may be distinctive to CNBD channel gating. First, traditional modular gating models assume voltage-independent coupling between voltage sensors and the pore, treating their interaction through allosteric factors that remain constant across membrane potentials (57). While this simplification successfully reproduces various types of voltage-dependent channel behaviors, emerging evidence now challenges the universality of voltage-invariant coupling. Recent structural studies of Kv and CNBD channels reveal that membrane voltage changes dynamically reconfigure the physical interface between voltage-sensing domains and the pore, raising the possibility that the voltage dependence of gating is determined, at least in part, by the coupling itself (17, 18, 43, 58). These findings collectively point to a possibility that VSD-PD couplings in VGICs inherently exhibit voltage sensitivity via voltage-dependent structural alterations, cooperative interactions, and state-dependent energetic landscapes that reshape transmission between voltage sensors and the pore. Furthermore, the voltage-independent closed-to-open transition appears to play an important role in determining the voltage-dependent gating polarity of CNBD channels. The bipolar U-shaped G-V requires a large L, indicating these mutations or chimeric reassembly facilitate the spontaneous channel opening. This is likely caused by the nondomain-swapped subunit arrangement of CNBD channels, characterized by the proximity between the VSD and PD, and a short S4-S5 linker. As a result, the intrinsic tendency of the pore opening is sensitive to even subtle changes in the VSD/PD interface. In contrast, to achieve proper reversed gating in HCN channels, a small L is required. Finally, the correlation between a larger gating charge (z) and the bell-shaped G-V relationship suggests more extensive voltage sensor movement. When the C-terminal end of spHCN channels was replaced with flexible alanine residues— particularly in the context of the unusual C-linker position in sea urchin sperm HCN channels—the S4 helix may have been able to move further downward (17, 31). Similarly, removing the PAS domain in Eag channels could enhance the extent of S4 movement (34). These findings suggest that the diversity in gating polarity among CNBD channels is directly influenced by the regulatory roles of their intracellular domains (49).

Fluorescence-based detection of transmembrane helix movements presents significant challenges. While hetero-FRET strategies, particularly using transition metal FRET, can be readily engineered to monitor large-scale vertical displacements (16, 17, 45, 59), detecting subtle lateral movements of helices within the membrane plane remains more difficult. Anisotropy-based homo-FRET offers an efficient alternative, as it relies on energy transfer between identical fluorophores, making it highly sensitive to their proximity in the membrane. This approach is particularly suited for studying concerted lateral motions, such as those in channel proteins with a 4-fold symmetric arrangement. However, accurate interpretation of anisotropy data depends on careful control of both fluorophore labeling efficiency and the selection of labeling sites. Future work should integrate site-specific fluorophore incorporation with patch-clamp electrophysiology to enable precise voltage control, providing a better understanding of the lateral movements and coupling between S4, S5, and S6 helices in CNBD channels.

## Methods

### Molecular biology, reagents and fluorescence imaging

The tsA-201 cell, a variant of human embryonic kidney (HEK) cells, was sourced from Sigma-Aldrich (St. Louis, MO). These cells were authenticated via STR profiling and stored in liquid nitrogen. For each new batch, the cells were cultured in Dulbecco’s Modified Eagle’s Medium (DMEM; Gibco), supplemented with 10% fetal bovine serum (FBS; Gibco) and 1% penicillin/streptomycin (Gibco). Cultures were maintained in a humidified incubator at 37°C with 5% CO_2_ in tissue culture dishes (CellTreat, Pepperell, MA). Transfection was carried out on tsA-201 cells at 70%-90% confluency using the Lipofectamine 3000 Kit (Invitrogen, Carlsbad, CA, #L30000080), following established protocols (60, 61). The hHCN4 and hERG1-S620T cDNAs were synthesized by VectorBuilder (Chicago, IL) and their expression was driven by a CMV promoter. S620T in hERG eliminates rapid C-type inactivation (62, 63) for isolating channel activation. Point mutations were induced using the QuikChange II XL site-directed mutagenesis kit (Agilent Technologies, Cedar Creek, TX). Verification of the introduced stop codons within the mutant plasmid was performed through DNA sequencing provided by Genewiz (Azenta Life Sciences, Burlington, MA).

The incorporation of L-Anap was conducted with the pANAP plasmid (Addgene plasmid #48696), the channel construct with the specific site mutated to *amber* stop codon (64), and with 20 μM L-Anap methyl ester (AsisChem, Waltham, MA) in the cell culture. To enhance nonsense suppression, a human dominant-negative release factor eRF1 (E55D) (VectorBuilder, Chicago, IL) was co-transfected (65). An eYFP was fused to the C-terminal end of the channel construct, as a validation of full-length expression of channels. Cells that had substantial membrane-localized YFP fluorescence were selected for imaging. Before imaging, cells were cultured in the L-Anap-free dPBS buffer for at least 10 mins, for gently washing away unincorporated L-Anap. To excite L-Anap, a 375 nm pulsed diode laser (ISS Inc, Champaign, IL) was employed. The emitted fluorescence from L-Anap was captured using a 447/60 nm band-pass emission filter.

An Olympus IX73 inverted microscope (Evident Scientific, Waltham, MA) and a laser-scanning confocal Q2 system (ISS Inc, Champaign, IL) were used for live-cell fluorescence anisotropy imaging. Anisotropy measurements were performed using a polarization module integrated with the confocal microscope, enabling simultaneous acquisition. The system included a halfwave plate and a linear polarizer assembly at the laser entrance, both of which were rotatable with 1-degree precision over 360 degrees to define the excitation polarization state. Emission fluorescence was separated into parallel (I_^∥^_) and perpendicular (I_⊥_) components relative to the excitation polarization using a polarization beam splitter assembly, allowing for precise anisotropy measurements. Anisotropy was determined using a software package in VistaVision (ISS Inc) that acquired fluorescence intensity in the two detection channels, with background subtraction applied to correct for autofluorescence and cytosolic fluorescence. The fluorescence anisotropy (r) at the plasma membrane was calculated using the equation: r = (I_∥_− I_⊥_) / (_∥_ + 2*I_⊥_) (52, 55). The data were presented as the mean ± SEM of n independent cells. Statistical significance was determined using the two-sided student’s t test.

### The modular allosteric model for CNBD channels

The values of the free parameters were estimated by minimizing the sum of squared differences between the targeted data and the simulated data using the Solver add-in in Microsoft Excel. The goodness of fit was quantified using the coefficient of determination (R^2^) and the root mean square error (RMSE). Data from the model were plotted using Igor Pro 9 (Wavemetrics, Portland, OR). The modular gating model consists of 4 independent VSDs per channel, each with two coupling factors to the PD: positive coupling factor *D*, and negative coupling factor *H*:

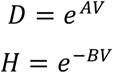

Where *A* and *B* are coupling strength parameters. *D* increases with depolarization and *H* increases with hyperpolarization. When *A* or *B* equals 0, then no coupling for *H* or *D*, or in other words, the S4 movement is not coupled to the pore. The charge associated with couplings were calculated using Z_coupling_ = Z_A_ + Z_B_ = (A + B)*RT/F.

The equilibrium constant for VSD activation is:

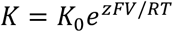

Where *K*_*0*_ is a constant 1.26E+6, z is gating charge, *F* is Faraday’s constant, *V* is membrane potential, *R* is the gas constant, and *T* is temperature in Kelvin (295 K).

The channel opening (*P*_*o*_) for the 8-state model was calculated based on the scheme in Fig. 2B:

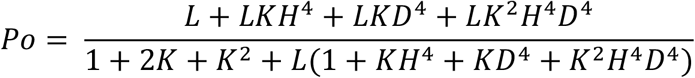

Where *L* is the intrinsic pore opening equilibrium. Thus, there are 4 free parameters in total in this model to compute *Po*-*V* relationships: *A, B, L*, and *z*. For unipolar gating behaviors, 3 free parameters including one coupling factor (*A* or *B, L* and *z*) are sufficient for the models. The 1 + *K* + *K*^*2*^ terms arise from a simplified equilibrium occupancy of independent voltage sensors and do not imply obligatory sequential activation steps of the VSD. For simplicity in data presentation, maximum *Po* for HCN and hERG was normalized and modeled to be 1 and shown as G/G_max_.

In addition, the channel opening (*P*_*o*_) for the 4-state model was similarly calculated based on the scheme of Fig. 3A:

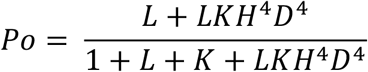

And the channel opening (*P*_*o*_) for the 6-state model was calculated based on the scheme of Fig. 3C:

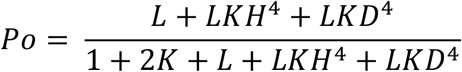

## Acknowledgements

We thank Natalie L. Macchi, Erin N. Lessie and Lucas J. Handlin for technical support. This research is supported by United States of America National Institutes of Health R35GM154778 and R56HL169176 grants, and by the Doisy Fund of the Department of Biochemistry and Molecular Biology at Saint Louis University School of Medicine.

## References

1. B. Hille, Ion Channels of Excitable Membranes (Sinauer Associates, ed. 3rd 2001).

2. Z. M. James, W. N. Zagotta, Structural insights into the mechanisms of CNBD channel function. J Gen Physiol 150, 225–244 (2018).

3. D. DiFrancesco, Characterization of single pacemaker channels in cardiac sino-atrial node cells. Nature 324, 470–473 (1986).

4. R. B. Robinson, S. A. Siegelbaum, Hyperpolarization-activated cation currents: from molecules to physiological function. Annu Rev Physiol 65, 453–480 (2003).

5. K. B. Craven, W. N. Zagotta, CNG and HCN channels: two peas, one pod. Annu Rev Physiol 68, 375–401 (2006).

6. D. DiFrancesco, Funny channel gene mutations associated with arrhythmias. J Physiol 591, 4117–4124 (2013).

7. W. A. Catterall, G. Wisedchaisri, N. Zheng, The chemical basis for electrical signaling. Nat Chem Biol 13, 455–463 (2017).

8. H. A. DeBerg, P. S. Brzovic, G. E. Flynn, W. N. Zagotta, S. Stoll, Structure and Energetics of Allosteric Regulation of HCN2 Ion Channels by Cyclic Nucleotides. J Biol Chem 291, 371–381 (2016).

9. J. Rheinberger, X. Gao, P. A. Schmidpeter, C. M. Nimigean, Ligand discrimination and gating in cyclic nucleotide-gated ion channels from apo and partial agonist-bound cryo-EM structures. Elife 7 (2018).

10. D. DiFrancesco, P. Tortora, Direct activation of cardiac pacemaker channels by intracellular cyclic AMP. Nature 351, 145–147 (1991).

11. M. C. Sanguinetti, C. Jiang, M. E. Curran, M. T. Keating, A mechanistic link between an inherited and an acquired cardiac arrhythmia: HERG encodes the IKr potassium channel. Cell 81, 299–307 (1995).

12. D. K. Jones et al., hERG 1b is critical for human cardiac repolarization. Proc Natl Acad Sci U S A 111, 18073–18077 (2014).

13. M. C. Trudeau, J. W. Warmke, B. Ganetzky, G. A. Robertson, HERG, a human inward rectifier in the voltage-gated potassium channel family. Science 269, 92–95 (1995).

14. E. C. Baker et al., Functional Characterization of Cnidarian HCN Channels Points to an Early Evolution of I_h_. PLoS One 10, e0142730 (2015).

15. S. Vemana, S. Pandey, H. P. Larsson, S4 movement in a mammalian HCN channel. J Gen Physiol 123, 21–32 (2004).

16. G. Dai, T. K. Aman, F. DiMaio, W. N. Zagotta, The HCN channel voltage sensor undergoes a large downward motion during hyperpolarization. Nat Struct Mol Biol 26, 686–694 (2019).

17. G. Dai, T. K. Aman, F. DiMaio, W. N. Zagotta, Electromechanical coupling mechanism for activation and inactivation of an HCN channel. Nat Commun 12, 2802 (2021).

18. R. Ramentol, M. E. Perez, H. P. Larsson, Gating mechanism of hyperpolarization-activated HCN pacemaker channels. Nat Commun 11, 1419 (2020).

19. G. A. Robertson, J. H. Morais-Cabral, hERG Function in Light of Structure. Biophys J 118, 790–797 (2020).

20. H. R. Guy, S. R. Durell, J. Warmke, R. Drysdale, B. Ganetzky, Similarities in amino acid sequences of Drosophila eag and cyclic nucleotide-gated channels. Science 254, 730 (1991).

21. G. A. Robertson, J. M. Warmke, B. Ganetzky, Potassium currents expressed from Drosophila and mouse eag cDNAs in Xenopus oocytes. Neuropharmacology 35, 841–850 (1996).

22. J. Warmke, R. Drysdale, B. Ganetzky, A distinct potassium channel polypeptide encoded by the Drosophila eag locus. Science 252, 1560–1562 (1991).

23. J. W. Warmke, B. Ganetzky, A family of potassium channel genes related to eag in Drosophila and mammals. Proc Natl Acad Sci U S A 91, 3438–3442 (1994).

24. W. Wang, R. MacKinnon, Cryo-EM Structure of the Open Human Ether-a-go-go-Related K+ Channel hERG. Cell 169, 422–430 e410 (2017).

25. J. R. Whicher, R. MacKinnon, Structure of the voltage-gated K+ channel EAG1 reveals an alternative voltage sensing mechanism. Science 353, 664–669 (2016).

26. T. I. Brelidze, A. E. Carlson, B. Sankaran, W. N. Zagotta, Structure of the carboxy-terminal region of a KCNH channel. Nature 481, 530–533 (2012).

27. T. I. Brelidze, E. C. Gianulis, F. DiMaio, M. C. Trudeau, W. N. Zagotta, Structure of the C-terminal region of an ERG channel and functional implications. Proc Natl Acad Sci U S A 110, 11648–11653 (2013).

28. T. I. Brelidze, A. E. Carlson, W. N. Zagotta, Absence of direct cyclic nucleotide modulation of mEAG1 and hERG1 channels revealed with fluorescence and electrophysiological methods. J Biol Chem 284, 27989–27997 (2009).

29. Y. Zhao, M. P. Goldschen-Ohm, J. H. Morais-Cabral, B. Chanda, G. A. Robertson, The intrinsically liganded cyclic nucleotide-binding homology domain promotes KCNH channel activation. J Gen Physiol 149, 249–260 (2017).

30. D. L. Prole, G. Yellen, Reversal of HCN channel voltage dependence via bridging of the S4-S5 linker and Post-S6. J Gen Physiol 128, 273–282 (2006).

31. G. E. Flynn, W. N. Zagotta, Insights into the molecular mechanism for hyperpolarization-dependent activation of HCN channels. Proc Natl Acad Sci U S A 115, E8086–E8095 (2018).

32. M. Tristani-Firouzi, J. Chen, M. C. Sanguinetti, Interactions between S4-S5 linker and S6 transmembrane domain modulate gating of HERG K+ channels. J Biol Chem 277, 18994–19000 (2002).

33. J. Cowgill et al., Bipolar switching by HCN voltage sensor underlies hyperpolarization activation. Proc Natl Acad Sci U S A 10.1073/pnas.1816724116 (2018).

34. H. Terlau, S. H. Heinemann, W. Stuhmer, O. Pongs, J. Ludwig, Amino terminal-dependent gating of the potassium channel rat eag is compensated by a mutation in the S4 segment. J Physiol 502 (Pt 3), 537–543 (1997).

35. C. H. Lee, R. MacKinnon, Structures of the Human HCN1 Hyperpolarization-Activated Channel. Cell 168, 111–120 e111 (2017).

36. S. B. Long, E. B. Campbell, R. Mackinnon, Voltage sensor of Kv1.2: structural basis of electromechanical coupling. Science 309, 903–908 (2005).

37. J. R. Whicher, R. MacKinnon, Regulation of Eag1 gating by its intracellular domains. Elife 8 (2019).

38. J. Xue, Y. Han, W. Zeng, Y. Wang, Y. Jiang, Structural mechanisms of gating and selectivity of human rod CNGA1 channel. Neuron 10.1016/j.neuron.2021.02.007 (2021).

39. M. Li et al., Structure of a eukaryotic cyclic-nucleotide-gated channel. Nature 542, 60–65 (2017).

40. Z. M. James et al., CryoEM structure of a prokaryotic cyclic nucleotide-gated ion channel. Proc Natl Acad Sci U S A 114, 4430–4435 (2017).

41. E. Lorinczi et al., Voltage-dependent gating of KCNH potassium channels lacking a covalent link between voltage-sensing and pore domains. Nat Commun 6, 6672 (2015).

42. A. P. Tomczak et al., A new mechanism of voltage-dependent gating exposed by KV10.1 channels interrupted between voltage sensor and pore. J Gen Physiol 149, 577–593 (2017).

43. X. Wu, K. P. Cunningham, R. Ramentol, M. E. Perez, H. P. Larsson, Similar voltage-sensor movement in spHCN channels can cause closing, opening, or inactivation. J Gen Physiol 155 (2023).

44. A. N. Chan et al., Voltage sensor conformations induced by LQTS-associated mutations in hERG potassium channels. Nat Commun 16, 7126 (2025).

45. A. N. Chan et al., Voltage Sensor Conformations Induced by LQTS-associated Mutations in hERG Potassium Channels. bioRxiv 10.1101/2024.05.17.594747 (2025).

46. M. Zhang, J. Liu, G. N. Tseng, Gating charges in the activation and inactivation processes of the HERG channel. J Gen Physiol 124, 703–718 (2004).

47. C. H. Lee, R. MacKinnon, Voltage Sensor Movements during Hyperpolarization in the HCN Channel. Cell 179, 1582–1589 e1587 (2019).

48. K. Ryu, G. Kasuya, K. Nakajo, Extracellular salt bridge networks around S4 implicated in HCN channel gating and heart disease. Proc Natl Acad Sci U S A 122, e2502136122 (2025).

49. J. H. Morais-Cabral, G. A. Robertson, The enigmatic cytoplasmic regions of KCNH channels. J Mol Biol 427, 67–76 (2015).

50. V. Burtscher et al., Structural basis for hyperpolarization-dependent opening of human HCN1 channel. Nat Commun 15, 5216 (2024).

51. A. Chatterjee, J. Guo, H. S. Lee, P. G. Schultz, A genetically encoded fluorescent probe in mammalian cells. J Am Chem Soc 135, 12540–12543 (2013).

52. G. Dai, Z. M. James, W. N. Zagotta, Dynamic rearrangement of the intrinsic ligand regulates KCNH potassium channels. J Gen Physiol 150, 625–635 (2018).

53. T. Kalstrup, R. Blunck, Dynamics of internal pore opening in Kv channels probed by a fluorescent unnatural amino acid. Proc Natl Acad Sci U S A 110, 8272–8277 (2013).

54. A. N. Bader et al., Homo-FRET imaging as a tool to quantify protein and lipid clustering. Chemphyschem 12, 475–483 (2011).

55. A. Joshi et al., Intermolecular energy migration via homoFRET captures the modulation in the material property of phase-separated biomolecular condensates. Nat Commun 15, 9215 (2024).

56. N. Decher, J. Chen, M. C. Sanguinetti, Voltage-dependent gating of hyperpolarization-activated, cyclic nucleotide-gated pacemaker channels: molecular coupling between the S4-S5 and C-linkers. J Biol Chem 279, 13859–13865 (2004).

57. F. T. Horrigan, R. W. Aldrich, Coupling between voltage sensor activation, Ca2+ binding and channel opening in large conductance BK potassium channels. J Gen Physiol 120, 267–305 (2002).

58. A. I. Fernandez-Marino et al., Inactivation of the Kv2.1 channel through electromechanical coupling. Nature 622, 410–417 (2023).

59. L. J. Handlin, G. Dai, Direct regulation of the voltage sensor of HCN channels by membrane lipid compartmentalization. Nat Commun 14, 6595 (2023).

60. G. Dai, Neuronal KCNQ2/3 channels are recruited to lipid raft microdomains by palmitoylation of BACE1. J Gen Physiol 154 (2022).

61. G. Dai, H. Yu, M. Kruse, A. Traynor-Kaplan, B. Hille, Osmoregulatory inositol transporter SMIT1 modulates electrical activity by adjusting PI(4,5)P2 levels. Proc Natl Acad Sci U S A 113, E3290–3299 (2016).

62. I. M. Herzberg, M. C. Trudeau, G. A. Robertson, Transfer of rapid inactivation and sensitivity to the class III antiarrhythmic drug E-4031 from HERG to M-eag channels. J Physiol 511 (Pt 1), 3–14 (1998).

63. S. J. Codding, M. C. Trudeau, The hERG potassium channel intrinsic ligand regulates N- and C-terminal interactions and channel closure. J Gen Physiol 151, 478–488 (2019).

64. S. E. Gordon, M. Munari, W. N. Zagotta, Visualizing conformational dynamics of proteins in solution and at the cell membrane. Elife 7 (2018).

65. W. H. Schmied, S. J. Elsasser, C. Uttamapinant, J. W. Chin, Efficient multisite unnatural amino acid incorporation in mammalian cells via optimized pyrrolysyl tRNA synthetase/tRNA expression and engineered eRF1. J Am Chem Soc 136, 15577–15583 (2014).

